# Capturing in saliva real time emotions induced by a fragrance: an objective emotional assessment based on multiplex molecular biomarker profiles

**DOI:** 10.1101/2025.09.16.676485

**Authors:** Laurence Molina, Francisco Santos Schneider, Malik Kahli, Alimata Ouedraogo, Mellis Alali, Agnès Almosnino, Julie Baptiste, Jeremy Boulestreau, Martin Davy, Juliette Houot, Telma Mountou, Marine Quenot, Elodie Simphor, Victor Petit, Franck Molina

## Abstract

This study introduces a non-invasive approach to objectively assess fragrance-induced emotions using multiplex salivary biomarker profiling. Traditional methods such as self-reports, physiological monitoring, or neuroimaging are often limited by subjectivity, invasiveness, or poor temporal resolution. Saliva offers a practical alternative, reflecting rapid neuroendocrine changes linked to emotional states.

We analyzed four key salivary biomarkers: cortisol (stress, HPA-axis activity), alpha-amylase (sympathetic activation), dehydroepiandrosterone (resilience), and oxytocin (social bonding, emotional regulation) to capture multidimensional emotional responses. Two clinical studies (N=30, N=63) and one consumer study (N=80) exposed healthy volunteers to six fragrances, with saliva collected before, 5 minutes after, and 20 minutes after olfactory stimulation. Subjective ratings of happiness, relaxation, confidence, and dynamism were also obtained via questionnaires. Rigorous analytical validation accounted for reproducibility, circadian variation and sample stability.

Biomarker patterns revealed fragrance-specific emotional profiles, with distinct subgroups of participants whose biomarker dynamics correlated with specific emotional states. Increased oxytocin and decreased cortisol consistently aligned with happiness and relaxation, whereas distinct biomarker combinations predicted confidence or dynamism. Classification and regression tree analysis demonstrated high sensitivity for detecting these profiles. Validation in an independent cohort (N=80) using an implicit association test confirmed concordance between molecular profiles and behavioral measures, underscoring the robustness of this method.

These findings establish salivary biomarker profiling as a reliable tool for decoding real-time emotional responses. Beyond scientific insights into affective neuroscience, this approach holds translational potential in personalized fragrance design, sensory marketing, and therapeutic applications for stress-related disorders. Expanding the biomarker panel and integrating molecular data with neuroimaging or autonomic measures could further elucidate the interplay between central olfactory processing and peripheral physiology.

## Introduction

Fragrances have long been recognized for their capacity to elicit strong emotional responses, ranging from pleasure and nostalgia to aversion and heightened alertness. Quantifying these effects presents a unique challenge: the act of measurement itself should not disrupt the emotional state being assessed. Traditional methods such as self-report questionnaires, behavioral observation, electroencephalography (EEG), functional magnetic resonance imaging (fMRI), or peripheral physiological monitoring (e.g., electrodermal activity, heart rate variability) have advanced the field but are often intrusive, limited in temporal resolution, or influenced by cognitive biases ^1-3^. At a biological level, olfactory stimuli evoke complex responses involving both central and peripheral systems. Signals from olfactory receptor neurons bypass the thalamus and project directly to primary olfactory cortex and limbic structures, notably the amygdala, hippocampus, and orbitofrontal cortex ^4,5^. These regions mediate both odor perception and emotional salience, engaging the hypothalamic– pituitary–adrenal (HPA) axis and the autonomic nervous system (ANS) to produce measurable changes in peripheral physiology ^6,7^. Despite these well-established links, there is currently no objective molecular method able to characterize emotional responses to olfactory stimulation in real time in peripheral fluids.

The amygdala responds rapidly to odor valence, even without conscious awareness ^5^. Pleasant and unpleasant odors elicit distinct activation patterns in the amygdala and orbitofrontal cortex ^8^, while hippocampal activity underlies the “Proust phenomenon” of odor-evoked autobiographical memory ^9^. These central processes are tightly coupled to peripheral outputs: stress-related odors can increase alpha amylase release ^10^, pleasant odors can reduce cortisol levels ^7^ and pain unpleasantness ^11^. Moreover, certain odorants modulate sympathetic activity within seconds ^6^. Such pathways highlight the integration between central detection, valence coding, and peripheral effectors (including hormones, neuropeptides, and enzymes), many of which can be detected in accessible biofluids such as saliva.

Existing tools for assessing emotion include self-report, physiological monitoring, and neuroimaging. Self-report measures capture the subjective experience but are vulnerable to introspection errors and social desirability bias ^12^. Physiological indices such as heart rate or skin conductance provide objective autonomic data yet lack the molecular specificity needed to dissect the biochemical mechanisms of emotion ^13^. Neuroimaging and electrophysiological techniques offer high neural resolution but are costly, stationary, and often intrusive ^14,15^. While valuable, these methods do not deliver a direct molecular readout of the peripheral physiological signature of emotion, particularly in response to sensory stimulation, and many are unsuitable for real-time or repeated use in naturalistic settings ^16^.

A complementary approach is to measure emotions through biofluids that capture rapid physiological dynamics. Such a matrix must have a rapid turnover and be sensitive enough to capture moment-to-moment changes, must reflect CNS-regulated neuroendocrine activity, and allow repeated non-invasive sampling without medical staff ^17-20^. Saliva fulfils all these criteria. It contains hormones, neuropeptides, enzymes, and metabolites whose concentrations vary over minutes to hours and are influenced by CNS and autonomic activity ^21,22^. Its proximity to the CNS and the permeability of the blood–brain barrier allows it to reflect acute neuroendocrine changes, while its ease of collection minimizes emotional disturbance during sampling.

A growing body of empirical research links salivary biomarkers to emotional and stress responses, each reflecting a distinct physiological pathway. Cortisol (CRT), the end product of HPA-axis activation, typically rises within minutes after stress onset and modulates emotional memory and vigilance ^2,23^. Odor-induced stressors have been shown to modulate salivary cortisol ^7^. Salivary alpha-amylase (sAA), a surrogate marker of sympathetic activation, rises within minutes of emotional arousal, offering high temporal resolution for detecting acute changes in stress and alertness ^18,24^. Dehydroepiandrosterone (DHEA), produced by the adrenal cortex, has neuroprotective and anti-glucocorticoid effects and has been associated with emotional resilience ^25,26^. Oxytocin (OXT), a neuropeptide involved in social bonding and affect regulation, can modulate amygdala reactivity to emotional stimuli ^27^, and changes in salivary OXT have been reported following social and sensory stimulation ^28,29^. Together, these biomarkers (BMs) enable multidimensional profiling of emotional states, spanning stress, arousal, resilience, and social-affiliative processes.

Few studies have combined multiple salivary biomarkers to profile emotional responses to odors. Most have focused on stress paradigms ^3,6^, leaving a gap in understanding how positive, neutral, or socially relevant odors are represented in peripheral molecular signatures. Integrating multiple biomarkers allows differentiation between emotional states that share similar arousal but differ in valence or in their relevance to social contexts.

In this work, we showed that saliva could capture the peripheral molecular signature of emotion in real time during olfactory stimulation. Simultaneous measurement of cortisol, sAA, DHEA, and OXT can reflect a substantial portion of the complexity of human emotional responses. We further hypothesize that distinct emotional states (or simplified valence/arousal profiles) can be mapped to reproducible biomarker variation patterns. This approach offers a non-invasive, scalable method for objective emotional assessment in both research and applied contexts.

## Materials and Methods

### Participants

All procedures were performed in accordance with applicable guidelines and regulations and informed consent was obtained from all participants prior to their inclusion in the study. The study was approved by a French national ethic committee (CPP-SUD EST I) on October 10, 2022 (22.03698.000189). The panel consisted of healthy volunteers aged 18–60 years, who were either naïve or trained in olfactory stimulation.

### Saliva collection and storage

Participants were required to refrain from eating, drinking or smoking for at least 30 min prior to saliva collection to minimize contamination risks. Unstimulated saliva samples were collected by passive drooling into sterile polystyrene tubes, with a minimum volume of 2 mL obtained for each sample. Visual inspection confirmed the absence of blood contamination. Immediately after collection, the samples were placed on ice and processed within the same day of collection. After treatment, samples were used immediately or stored at −80°C.

### Clinical studies

We designed and conducted two clinical studies and a consumer test, each involving a separate group of volunteer subjects (N=30, N=63, N=80, respectively). For the clinical research studies (N=30 and N=63), participants smelled one fragrance per day, in the morning and the order of the fragrances were randomized for all participants. Saliva samples were collected at three different times during the panel: 5 minutes before stimulation by a specific olfactory fragrance (S1), 5 minutes after stimulation (S2), and 20 minutes after stimulation (S3) (Fig. 1). Six different fragrances were studied (two fragrances in the first study and four in the second). Participants completed a questionnaire about their emotional responses immediately after stimulation, while continuing to scent the fragrance for 5 minutes, just before the S2 sample was collected.

**Fig. 1:**
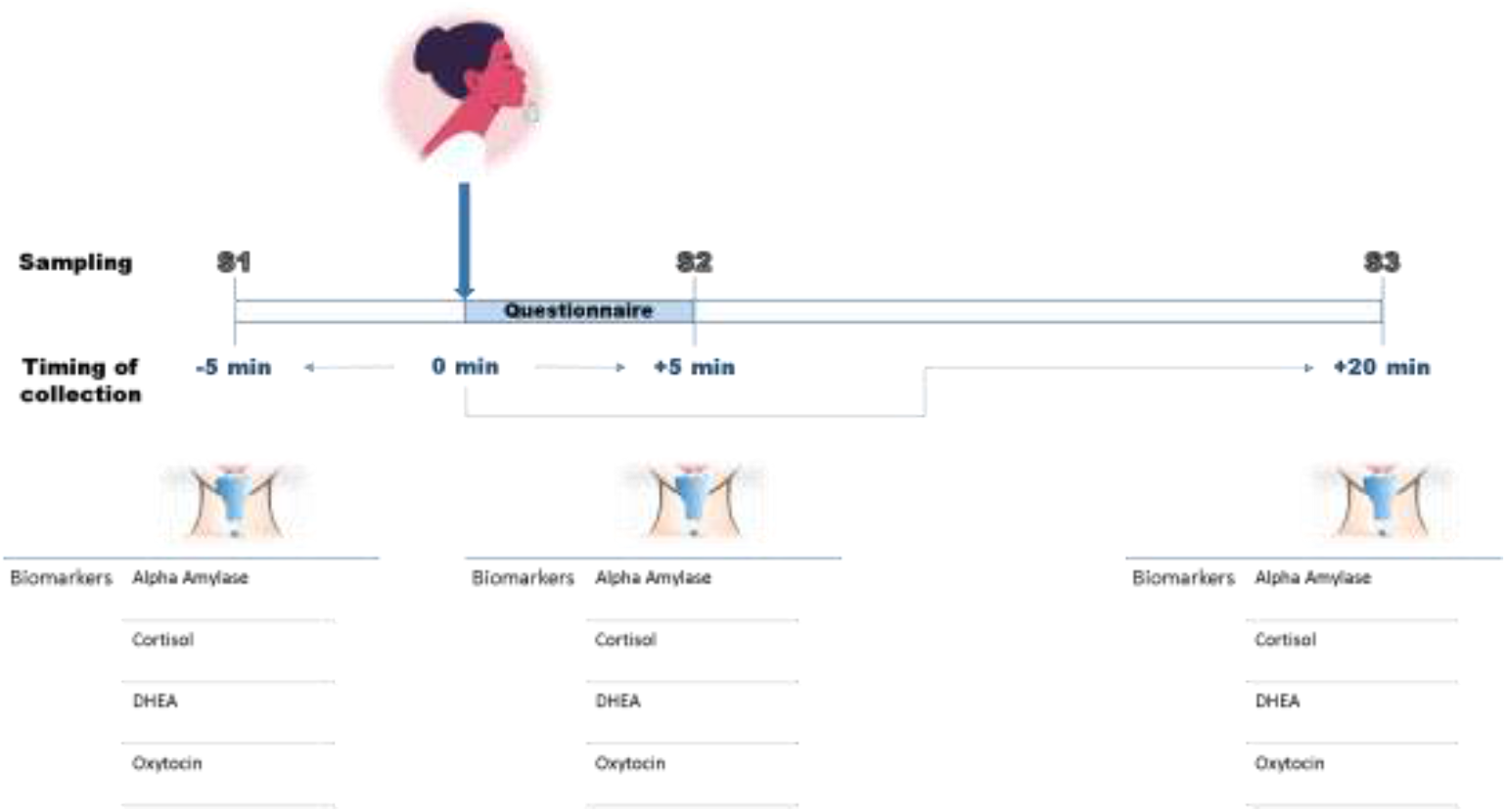
Design of the clinical research studies. Each participant scented two or four different fragrances depending on the clinical study (one fragrance per day of participation). The participation consisted in 1) a 5-minute sit-down-rest before the first sampling of saliva (5 minutes before olfactory stimulation), 2) scent of the fragrance and response to the questionnaire, 3) a second saliva sampling five minutes after scenting the fragrance and finally 4) a third saliva sampling 20 minutes after smelling the fragrance.

For implicit test, the panelist (n=80) scent the same four fragrances as in the second clinical research study. Individual emotional evaluations were performed using a combination of questionnaire (six to nine emotions assessed) and implicit-association test.

### Biomarkers measurement in saliva

Salivary cortisol were assayed using a commercial competitive solid phase enzyme-linked immunosorbant assays (ELISA) purchased from Cayman Chemical (Michigan, USA). Salivary DHEA concentrations were determined using a commercial competitive ELISA purchased from Tecan (Hamburg, Germany). 50 µL of saliva samples for each assay were used for these kits. Salivary oxytocin levels were analyzed using a competitive ELISA purchased from Enzo Life Sciences (NY, USA) using 100 µL of saliva samples for each assay. Salivary alpha-amylase was measured using an enzymatic assay purchased from IBL International (Tecan, hamburg, Germany). 10 µL of saliva were used for each assay. Saliva samples were analysed immediately after collection or after storage at −80°C.

### Emotional assessment on panelists

For each of the olfactory stimulation, separate emotional assessments have been performed using a questionnaire. The questionnaire evaluated certain parameters related to fragrances, such as intensity and valuation, as well as the emotions aroused by the olfactory experience including “confident”, “sensual”, “happy”, “dynamised”, “relaxed’” and “comforted”. For the first clinical study, three additional emotions were assessed “elegant”, “addict” and “unique”. Participants classified their emotions’ feelings into levels ranging from 0 to 2 (0 lowest valence, 2 highest valence).

Independent implicit association test were done on separated set of panelists (N=80; 50% men; 50% women) using the same four fragrances. Participants who did not consider fragrances important and did not use them regularly were screened out. Subjects were instructed to press the space bar on the computer if the emotional attribute administered matched their emotional state when they felt the stimulus.

### Data analyses and statistics

Differences of quantitative variables were tested by Wilcoxon rank sum test, Kruskal-Wallis test or ANOVA. We corrected for multiple hypotheses with the Benjamini-Hochberg test. PCA was performed using the R function prcomp in the stats library ^30^. For the Linear Discriminant Analysis (LDA), a supervised method, a 2-fold cross-validation was used to compute accuracies, a mean of 100 repeats is given. LDA was performed with the R function lda of the MASS_7.3-60 library ^30^. K-means and hierarchical k-means were performed with the R function k-means in the stats library and hk-means in the factoextra_1.0.7 library, respectively. The hierarchical clustering step of the hk-means was carried out using the Euclidian distance to measure the distance between individuals and Ward’s minimum variance method combined to the squares Euclidian distance was used as a linkage method. Radar chart graphs were obtained with the function radarchart of the fmsb_0.7.6 R library. The Classification and Regression Trees (CART) algorithm was performed using discrete variables of S2/S1 ratios of biomarkers and executed using the rpart (version 4.1.24) and rpart.plot (version 3.1.2) libraries in R software. Concentration ratios (S2/S1) were discretized into three categories using the mean coefficient variation (10%): values < 0.9 were designated as D (Decrease), values between 0.9 and 1.1 as S (Stable), and values > 1.1 as I (Increase).

## Results

### 1. Reference testing and analytical validation of salivary biomarkers: stability, robustness, and circadian considerations

We first validated the measurement of four salivary biomarkers (cortisol, sAA, DHEA and OXT) by comparing two commercial assay kits for each molecule based on defined criteria (e.g., linearity, dynamic range, and assay reproducibility). Reference assays were selected based on performance in saliva, with intra- and inter-assay variations coefficients of variation (CVs) (Tables 1SM and 2SM). To assess sample stability, salivary biomarkers were tested after storage conditions under multiple temperature conditions (−20°C, 4°C, room temperature, 37°C, 42°C and 60°C for 48 hours) (Fig. 1SM). sAA, DHEA and cortisol remained relatively stable at −20°C, +4°C and RT, whereas oxytocin showed stability only at −20°C. Based on these results, all biomarkers were analyzed on the day of sample collection. The impact of circadian rhythm was assessed in samples collected throughout the day (Fig. 2SM). Salivary cortisol, DHEA and OXT concentrations were higher in the early morning (9 a.m.), declining progressively through the day, with a modest effect for OXT, while sAA activity presented the opposite pattern, markedly increasing it levels after midday. These findings are consistent with previously published data ^31,32^. Given that the olfactory stimulation during the clinical study last approximately 30 minutes and that short-term biomarker variations were minimal, approximately 6% over 30 min., we therefore decided to collect samples in the morning.

### 2. Assessing the emotional and physiological impact of olfactory stimuli

We next examined whether olfactory stimulation modulates salivary biomarkers and whether changes depend on the odorant, its support, or the test procedure itself. A proof-of-concept study was conducted including 30 participants, with repeated saliva sampling to capture biomarker kinetics following exposure to either a neutral fragrance carrier or a fragrance stimulus (Fig. 1). Across conditions, cortisol and DHEA were moderately correlated after both fragrance or carrier stimulation (Fig 3SM). In contrast, sAA showed a negative correlation with cortisol and DHEA, but only in response to the fragrance. OXT displayed a distinct pattern, diverging from the other BMs. These differences between fragrance and carrier were consistent across the three sampling time-points (S1, S2 and S3), supporting the value of retaining all four biomarkers to capture the multidimensional signature of emotional responses. Individual-level analyses revealed high inter-individual variability. More than half of the subjects exposed to the carrier reported little to no emotion in the questionnaire in parallel to minimal biomarker fluctuations (e.g., subject E-010 and E-024 for carrier fragrance; Fig 4SM). In contrast, only three subjects failed to report an emotional response to the fragrance. Several individuals (e.g., E004, E007, E010) showed marked biomarkers shifts within 5 min of odors exposure and also exhibited an highlighted emotional response. These data suggest that sensitivity to olfactory stimuli varies considerably across individuals and can be detected at the biomarker level. Unsupervised k-means clustering analysis of biomarker profile identified subgroups of participants whose biomarker dynamics aligned with distinct emotional responses (in particular for clusters 2, 4 and 5 (Fig. 5SM)). This exploratory analysis demonstrates that multivariate biomarker signatures can differentiate both stimulus-specific effects and inter-individual variability in fragrance-induced emotions.

### 3. Expanded cohort study using four different odors as stimuli

#### Biomarkers ratio performances

To extend our proof-of-concept findings, we conducted a larger study (N=63) using four different fragrances (J3-J6). Four participants provided two or three samples only, leaving a total N=59 participants with 4 sampling, resulting in 708 saliva samples across 236 participant-fragrance combinations and three sampling points (S1-S3; Fig. 1). The biomarkers were measured robustly, with mean coefficients of variation <9%.

Baseline concentrations (S1, before stimulation) revealed broad inter-individual variability (e.g. sAA: 79.1±57 U/mL; CORT: 2559.2±1534.6 pg/mL; DHEA: 745.3±460.3 pg/mL; OXT: 96.1±75.1 pg/mL). To account for this, responses were expressed as post-stimulation ratios relative to baseline (S2/S1, S3/S1).

Across the full cohort, weak correlations emerged between DHEA and cortisol, while the other biomarkers varies independently (Fig. 6SM). This independence confirms the complementary contribution of each BM to the molecular signature of emotions as observed in the first clinical study.

We next applied multiple analytical approaches to examine biomarker dynamics at population level: i) principal component analysis (PCA) (Fig. 7SM), ii) ANOVA (Fig. 8SM), iii) hierarchical k-means (hk-means) clustering of biomarker ratio profiles (Fig. 9SM and Fig 10SM) and iv) temporal clustering of baseline versus post-stimulation ratios (Fig. 11SM). Collectively, these analyses indicated associations between biomarker ratio profiles and reported emotional responses, although noise in the pooled dataset (all fragrance combined) limited the identification of clear BM-emotion pairings. Indeed, no significant differences between hk-means cluster were identified when emotional responses were discretized and assessed by Chi^2^ or PCA analyses (Fig. 12SM, Table 3SM).

#### Classification of panelist responses to specific fragrance

Because physiological and emotional responses to odors varied substantially across individuals and seems to depend on the stimuli, we next performed individual-level clustering analyses for each fragrance. K-means and hk-means were applied to S2/S1 biomarker ratios (Fig. 13SM). For example, stimulation with fragrance J3 generated five clusters (n = 21, 22, 7, 9 and 1), whereas fragrance J6 produced four clusters (n = 28, 25, 3 and 3). The nested architectures obtained with fragrance J5 and J6 were different from that obtained with J3 and J4 fragrances. To explore hierarchical structure, hk-means was repeated with varying thresholds and recursively applied to subclusters.

For each fragrance, biomarker distributions and emotional responses were systematically compared across clusters formed by hk-means (Fig. 2). Emotional responsiveness was classified as high (>66% reporting), moderate (>33% and <66%), or low (<33%). Using this framework, hk-means clusters exhibited distinct BM profiles that corresponded to specific emotion response profiles. For instance, clusters derived from fragrance J6 showed divergent combinations of happiness, relaxed, comforted, and dynamised, each associated with a unique salivary biomarker profile.

**Fig. 2:**
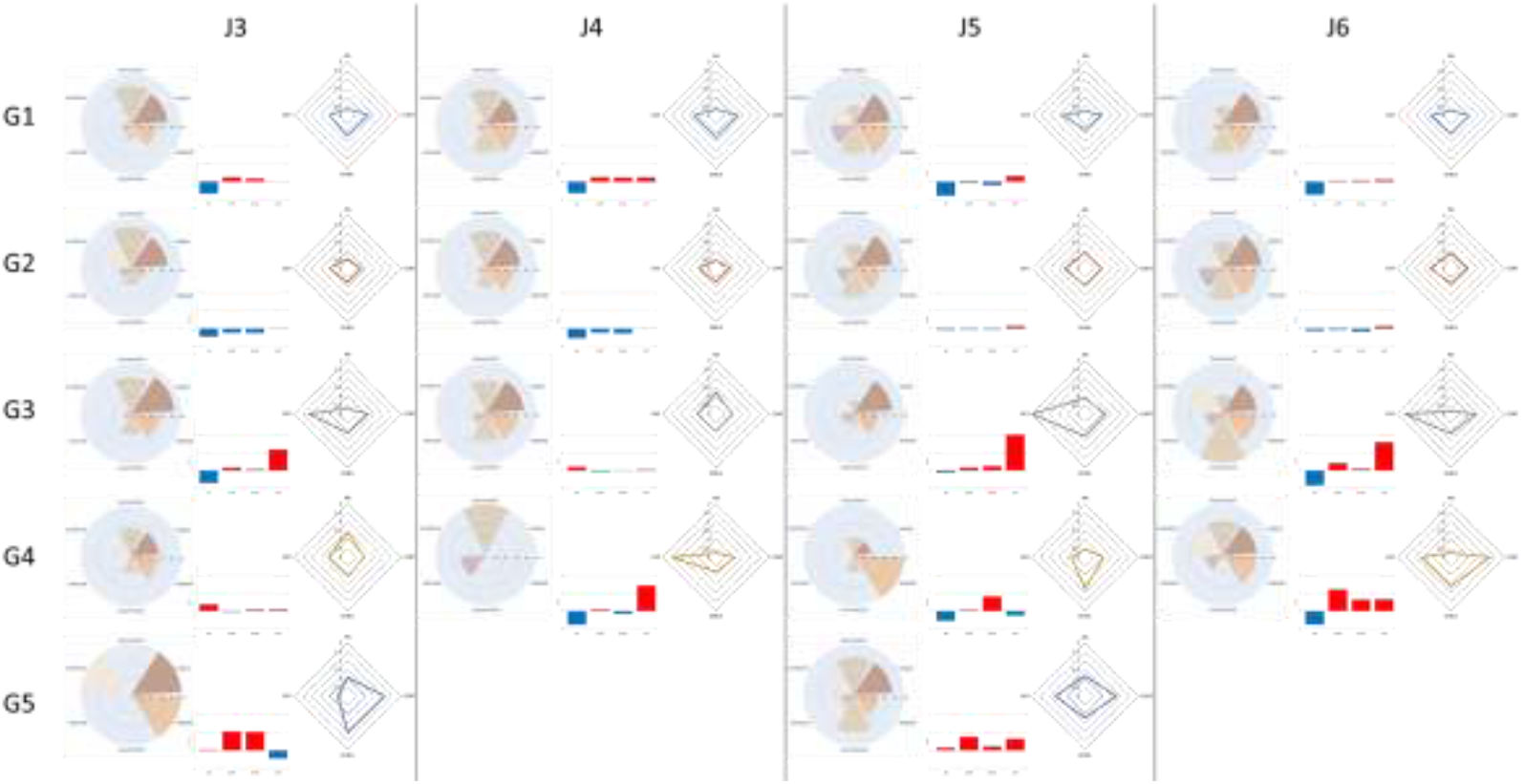
Emotional responses and salivary biomarker S2/S1 ratio profiles based on hk-means clusters by fragrance. For each fragrance (J3–J6), clusters are shown with emotional response profiles (left, radar plots; % participants per cluster) and log2 mean biomarker ratios (S2/S1) (center, histograms; right, spider plots).

Strikingly, similar biomarker–emotion associations emerged across different fragrances. Clusters J5G2 (Fragrance J5, group 2) and J6G2 both displayed modest decreases in sAA, cortisol and DHEA concomitant with increased OXT, and participants in these clusters reported highly comparable emotional response profiles characterized predominantly by high happiness and moderate relaxation. Similarly, clusters J3G2 and J4G2 were marked by pronounced decreases in sAA, cortisol and DHEA with low decreased/stable levels of oxytocin corresponding to emotional response profiles centered on happiness and dynamism.

Together, these findings show that olfactory stimulations elicits distinct molecular and emotional responses clusters within a population and that specific biomarker signatures can recur across different fragrances while maintaining consistent emotional associations. This suggests that clustering of salivary biomarkers provides a powerful framework for linking physiological responses to subjective emotional states.

#### Individual analyses of fragrance-induced molecular profiles reveal clear association with emotional response

At the individual level, salivary biomarker ratios profiles showed clear associations with reported emotions (Fig. 14SM). For instance, participant E-044 displayed low stable /increased/ levels of sAA and OXT, with marked decreases in cortisol and DHEA following exposure to fragrances J3 and J4, a profile associated with an absence of emotional responsiveness. In contrast, the same participant exhibited strong decreased in sAA, moderate/high increased levels of cortisol, DHEA and oxytocin when exposed to fragrances J5 and J6, corresponding to reports of moderate happiness and high sense of comfort and relaxation. Similar patterns were observed in other participants. For instance, E-078 displayed similar biomarker profiles for J3 and J5 characterized by elevated cortisol and DHEA with reduced oxytocin, aligning with relaxed emotional states and moderate happiness. Stimulation with J4 and J6 induced a different pattern, with decreases in all four biomarkers, that was associated with happiness, relaxation, and confidence. These cases illustrate how fragrance-specific shifts in biomarker ratios parallel the individual’s emotional experience (Fig. 14SM).

#### Identification of Biomarker Profiles linked to specific emotions

To reduce the signal-to-noise of biomarker ratios, data were discretized using the mean coefficient of variation (∼10%) as the threshold. Ratios were categorized as decreased (<0.9), stable (0.9–1.1), or increased (>1.1), generating 81 possible combinations of BMs states across 236 participants-fragrances exposures (Fig. 15SM).

Classification and Regression Trees (CART) were then used to predict emotional responses from biomarker profiles (Fig. 3). Using this approach, participants were classified into leaves indicating either a positive (green) or negative (blue) response to a given emotion.

**Fig. 3:**
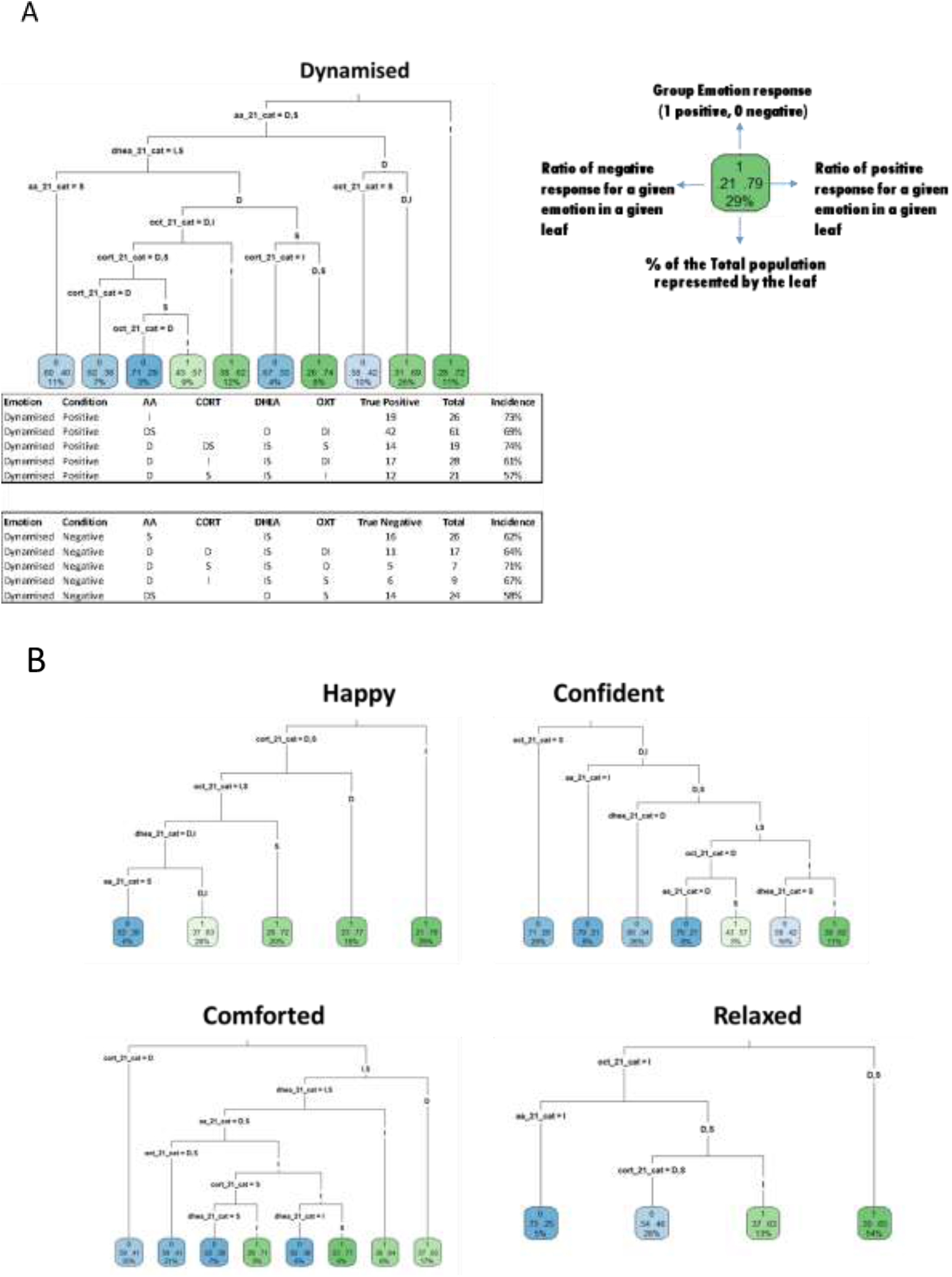
Classification of salivary biomarker ratio (S2/S1) profiles for emotion identification. The “Classification and Regression Trees” (CART) algorithm was performed using discrete variables of S2/S1 ratios of biomarkers. Participants were classified in the leaves according to whether they felt the emotion analyzed (positive: green) or not (negative: blue). The numbers and information inside leaves are explained in the top right-hand corner of the figure. **(A)** CART for the dynamised emotion. **(B)** CART for the emotions Happy, Confident, Comforted and Relaxed. The CART method was performed using the entire cohort data (236 unique combinations of individuals/fragrances).

For the emotion “dynamised” (Fig. 3A), CART analysis identified distinct biomarker configurations: 69% of subjects with a profile of decreased/stable sAA, decreased DHEA and variable OXT reported this emotion, representing 26% of the entire study population. Conversely, stable sAA combined with stable/increased DHEA was associated with the absence of this emotion. Comparable CART decision trees were generated for the emotions happy, confident, comforted and relaxed (Fig 3 B-D). For instance, 77% of people with stable DHEA, increased cortisol and oxytocin, and decreased/stable sAA reported feeling comforted.

Predictive performance varied by emotion (Table 4SM). Happy was identified with high sensitivity (97%) but low specificity (13%), whereas comforted and confident were predicted with high specificity (82% and 91%, respectively) but lower sensitivity (43% and 24%). These results indicate that discrete biomarker patterns capture emotion-specific molecular signatures, though predictive strength depends on the emotion assessed.

#### Emotional profiles of fragrances based on biomarkers profiles

The CART-derived biomarker patterns were then applied to characterize the emotional profiles induced by each fragrance (J3-J6). For each emotion, the proportion of participants with corresponding biomarker ratio profile was calculated (Fig. 4A). Each fragrance elicited a distinct emotional signature, confirming that olfactory stimulation can be differentiated at the molecular level.

**Fig. 4:**
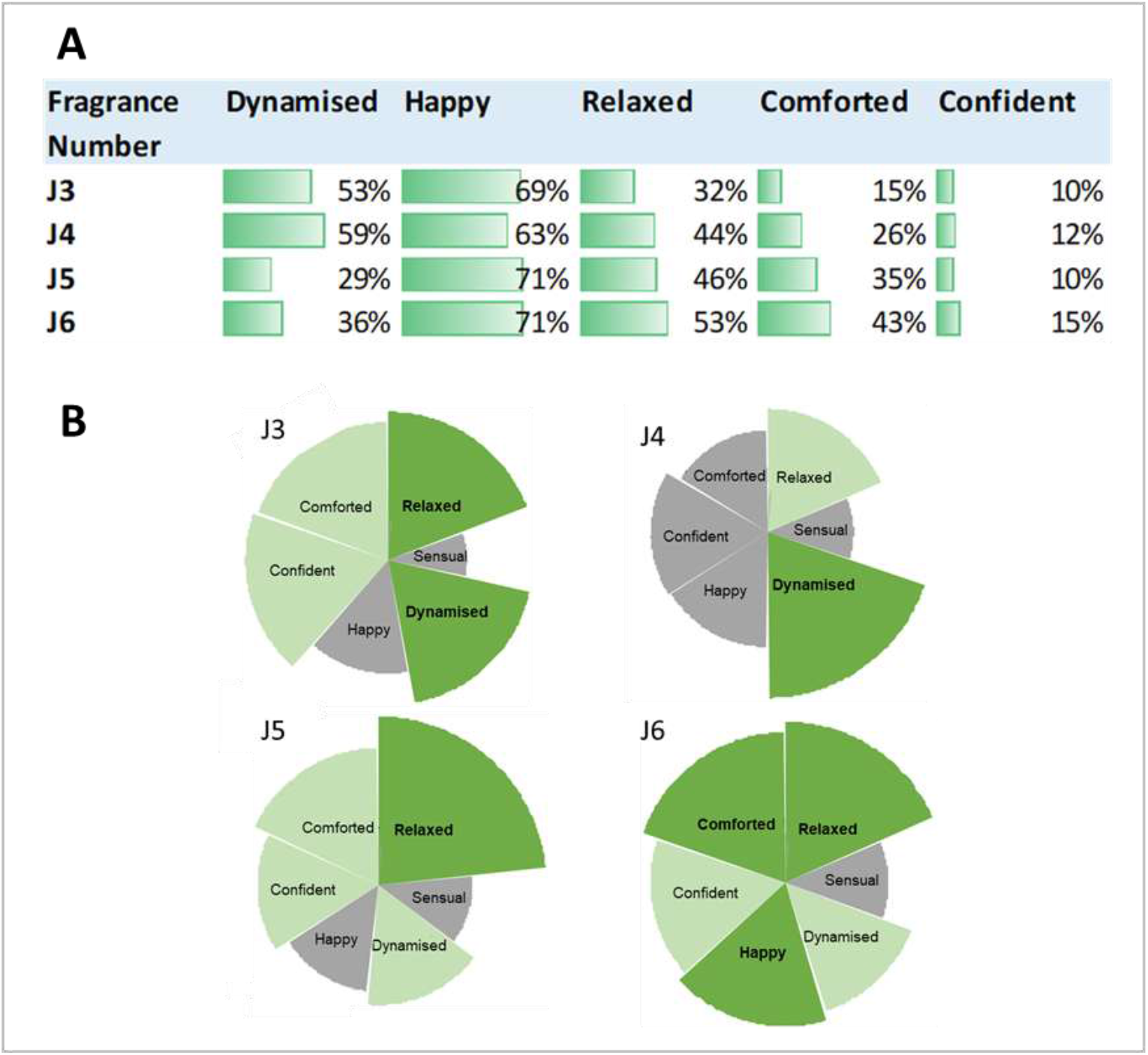
Biomarker-based emotional profiles induced by fragrances. **(A)** Salivary biomarker-based patterns associated to emotional responses were used to identify the emotional profiles of the tested fragrances (J3, J4, J5 and J6) in the cohort. Biomarkers-based emotional responses for each fragrance correspond to % of participants presenting the identified biomarker-based emotional profiles from the total number of participants that scented a given fragrance. Patterns used in this approach were those identified from the decision trees generated by CART performed on the entire cohort data (236 unique combinations of individuals/fragrances) and biomarkers’ ratios S2/S1 as discrete variables. **(B)** Implicit test-based emotional profiles induced by each fragrance. Two parameters were measured with the implicit test: the GO/NO GO percent (the number of people who validated the emotion for each fragrance) and the strength of the association (the speed of response to validate the emotion for each fragrance). The GO/NO GO percent was represented by the angle (emotions were presented 3 times during the test and the % represents the number of clicks for each emotion). The strength of association (speed of clicks) was represented by the size and the color. An association is considered high when rapid responses were given by the participant. Dark green : excellent (>80 percentile, emotion shown in bold); green : good (>50 percentile); grey : low (<50 percentile).

To validate these findings, an independent cohort (n = 80) completed an implicit association (IAT) test with the same four fragrances. Emotional profile derived from implicit measures (Fig. 4B) aligned closely with those inferred from biomarker data, supporting the robustness of the biomarker-based approach.

#### Role of individual biomarker in fragrance valuation

Finally, we examined whether individual biomarkers were associated with subjective fragrance valuation across the entire dataset (236 participant–fragrance combinations; Fig. 16SM). Participants who rated fragrances poorly (scores 5–6) exhibited lower sAA ratios and higher oxytocin ratios compared with those giving medium (3–4) or high (1–2) ratings. These findings suggest that sAA and oxytocin may contribute to the evaluative dimension of fragrance perception, linking molecular changes not only to emotional states but also to hedonic valuation.

## Discussion

This study provides proof-of-concept that salivary biomarkers can be objectively used to characterize emotional responses to olfactory stimulation in real time. Using validated assays with high reproducibility (mean CV <10% intra- and inter-assay for the main study), we quantified alpha-amylase, cortisol, dehydroepiandrosterone, and oxytocin in 888 saliva samples collected during controlled fragrance exposures. Our findings show that salivary molecular profiles can capture distinct physiological dimensions of emotion without the invasiveness or cognitive biases of self-report or neuroimaging ^19,33^.

### Multidimensional salivary profiling of olfactory emotion

Most previous olfactory–emotion research have measured only one biomarker, such as cortisol in odor-induced stress ^11^ or sAA changes during affective odor stimulation ^34^. Our integration of four functionally distinct biomarkers captures a broader physiological spectrum and enables discrimination between affective states that may share arousal intensity but differ in valence or affiliative meaning. Weak inter-correlations among cortisol, alpha-amylase, DHEA, and oxytocin underscore their complementary roles: cortisol reflects HPA-axis activation ^2^, sAA rises within minutes following sympathetic activation ^18,24^, DHEA modulates glucocorticoid effects and resilience ^25,26^ and oxytocin supports affiliative signaling ^28,29^. In stress physiology, combined profiles such as cortisol and sAA or DHEA:cortisol ratios have been used to stratify performance and anxiety responses ^35-37^, and here we extend this multidimensional approach to fragrance-evoked emotions.

### Individualized molecular signatures in olfactory emotion

Participants clustered into distinct biomarker-defined subgroups in response to the same fragrance, consistent with psychophysiological evidence of marked inter-individual variability in odor perception and affective responses ^6,38^. Neuroimaging studies have similarly shown that individuals with comparable subjective ratings can display divergent neural activation patterns to identical odor stimuli ^39^, suggesting trait-like differences in central–peripheral coupling. We observed reproducible biomarker–emotion patterns across different fragrances supporting the idea of stable molecular “emotional phenotypes”. This parallels the affective startle paradigm, in which eyeblink reflexes are potentiated during unpleasant stimuli and attenuated during pleasant ones, with stable inter-individual differences considered trait-like affective profiles ^40^. By analogy, the reproducible biomarker–emotion signatures observed here may reflect consistent molecular phenotypes of olfactory emotional processing.

### Machine learning for physiological emotion classification

Hierarchical k-means and CART decision trees analyses showed that discrete biomarker ratios can be linked to specific emotional states with transparent, rule-based classification. CART identified biomarker configurations that identify the presence or absence of emotions such as happiness, comfort, or confidence, with varying balances of sensitivity and specificity. This extends earlier work in affective computing, where physiological signals such as heart rate variability or skin conductance have classified emotional valence with >80% accuracy ^41^. While happiness was detected with high sensitivity but low specificity, comfort and confidence were classified with high specificity but lower sensitivity. These patterns highlight both the promise and the current limitations of molecular prediction. Importantly, the interpretability of decision trees is an advantage in biological contexts: rather than giving opaque predictions, they reveal decision rules that can be mechanistically tested. For instance, combinations of decreased sAA with increased oxytocin were consistently associated with comforted states, while stable sAA and increased DHEA predicted the absence of dynamised emotion. Such rules may generate hypotheses about the pathways connecting olfactory input to peripheral endocrine responses.

### Convergent validation with implicit measures

Construct validity was supported by the concordance between our biomarker-based profiles and an independent Implicit Association Test conducted in a separate cohort. Previous research has shown that implicit affective measures correlates with physiological stress markers, such as cortisol ^42^ and facial EMG ^43^. In this line, we found that implicit and biomarker-derived profiles converged for fragrance-evoked emotions. This cross-method consistency suggests that salivary biomarkers can provide an objective complement to behavioral measures in emotion research.

### Biomarkers and hedonic appraisal

We also observed associations between biomarkers and fragrance valuation. Lower alpha-amylase ratios and higher oxytocine ratios were associated to reduced liking, suggesting that these markers may encode evaluative as well as emotional dimensions of olfactory experience. While oxytocin is frequently associated with positive valence and prosociality ^28^, context-dependent negative effects have been reported ^44^, and sympathetic downregulation may reflect disengagement ^45^. Thus, biomarker responses appear to capture a complex interplay of arousal, affiliation, and appraisal, rather than mapping linearly to valence polarity.

### Implications for personalization

The identification of stable molecular subgroups points toward potential personalization of sensory interventions. In clinical contexts, peripheral endocrine profiles has been shown to predict psychotherapy outcomes in PTSD, with baseline cortisol metabolism, DHEA:cortisol ratios ^46,47^, and HPA-axis reactivity (via the cortisol awakening response), all stratifying treatment response ^48,49^. Analogously, biomarker-defined phenotypes could be used to tailor fragrance design or therapeutic olfactory applications to individual emotional sensitivity profiles. More broadly, this work illustrates how molecular signatures may enable individualized models of emotion that integrate subjective and biological readouts.

### Limitations and next steps

Our biomarker panel does not include catecholamines or inflammatory mediators, which also respond dynamically to affective stimuli ^50-52^. Other hormones or peptides present in saliva could be included as well. Assay variability remains a challenge, particularly for salivary oxytocin, and larger, more diverse cohorts are needed to confirm the stability of identified phenotypes. Finally, integrating molecular profiling with concurrent neural and autonomic measures will be essential to map the temporal dynamics linking central olfactory processing to peripheral molecular responses.

## Conclusion

This study demonstrates that multi-biomarker salivary profiling is a viable, reproducible, and non-invasive tool to assess olfactory-evoked emotions. By combining molecular data with machine learning and validating against implicit measures, we provide a robust, ecologically valid framework for decoding emotional responses. Multidimensional molecular profiling captures both shared and individual aspects of affective physiology, offering a bridge between central sensory processing and subjective experience. This approach has potential applications in affective neuroscience, sensory marketing, and personalized therapeutic interventions, and could be extended to other sensory domains where rapid, non-invasive emotion assessment is needed.

## Acknowledgments

The authors would like to thank Mrs Agnès Fournier and Caroline Dubourg and Mr Louis Aguadish and Loïc Bleuez, for supplying the fragrances and carrying out the implicit association tests. We sincerely thanks Alice René, responsible for the bioethic regulation at CNRS, for her valuable advice in the field of clinical research studies and the CNRS for being the promotor of the clinical studies. Statistical analyses were carried out with the technical facility StatABio BioCampus, UAR 3426, CNRS, Inserm, University Montpellier, Montpellier, France. We also thank all the study participants.

## Author contributions

Study concept and design: F.S.S, L.M, M.K and F.M. Acquisition, analysis, interpretation of data: All authors. Drafting of the manuscript and critical revision of the manuscript for important intellectual content: F.S.S, L.M, M.K and F.M. Statistical analysis: E.S., M.D. Funding acquisition: V.P. and F.M. All authors critically reviewed and edited the manuscript.

## Data availability statement

The authors declare that all relevant data have been provided within the manuscript and its supporting information files.

