## Supplemental materials for "Capturing in saliva real time emotions induced by a fragrance: an objective emotional assessment based on multiplex molecular biomarker profiles"

*2. SkillCell*
*1682 rue de la Valsière CS 61003, 34184 MONTPELLIER Cedex 4, France*

### Supplemental Materials

**Table 1SM: Intra-assay variability measured on salivary samples for each biomarkers with selected reference kits.** Values (%) represent the mean CVs calculated for the 8 salivary samples per assay and per operator. CV: coefficient of variation.

| | Mean intra-assay CV | | | | | |
|---|---|---|---|---|---|---|
| | Operator A | | | Operator B | | |
| **Biomarkers** | ***Assay 1*** | ***Assay 2*** | ***Assay 3*** | ***Assay 1*** | ***Assay 2*** | ***Assay 3*** |
| **A-Amylase** | 9% | 6% | 5% | 5% | 5% | 5% |
| **Cortisol** | 5% | 5% | 7% | 12% | 5% | 8% |
| **DHEA** | 8% | 10% | 9% | 6% | 12% | 9% |
| **Oxytocin** | 17% | 21% | 12% | 20% | 17% | 23% |

**Table 2SM: Inter-assay variability measured on the 8 salivary samples per biomarkers.** Values (%) represent averages of all CVs obtained for the same salivary sample tested six times (2 operators x 3 independent tests) with each biomarker. CV: coefficient of variation.

| | Mean inter-assay CV for each salivary sample | | | | | | | |
|---|---|---|---|---|---|---|---|---|
| **Salivary samples** | **S-12153** | **S-13152** | **S-22184** | **S-22516** | **S-38169** | **S-41131** | **S-65121** | **S-65821** |
| **A-Amylase** | 7% | 4% | 10% | 6% | 5% | 6% | 5% | 4% |
| **Cortisol** | 12% | 2% | 11% | 6% | 9% | 5% | 6% | 5% |
| **Salivary samples** | **S-11113** | **S-15211** | **S-18215** | **S-21510** | **S-22147** | **S-22184** | **S-41131** | **S-65121** |
| **DHEA** | 8% | 9% | 13% | 9% | 8% | 8% | 8% | 8% |
| **Ocytocine** | 22% | 12% | 21% | 13% | 14% | 20% | 15% | 29% |

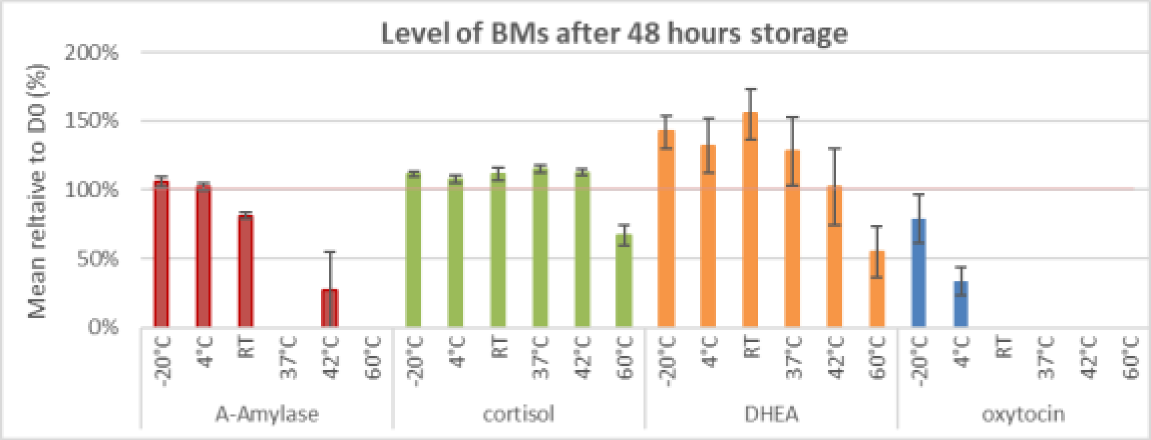


**Fig. 1SM: Stability of biomarkers as a function of storage temperature for 48 hours.** Results represent averages of measured biomarker values in human salivary samples (N=4). Results are expressed in % relative to the concentration measured at Day 0 (D0), which represents 100%. D0 assays were performed on fresh saliva. Mean CVs are represented by bars. RT: room temperature; CV: coefficient of variation.

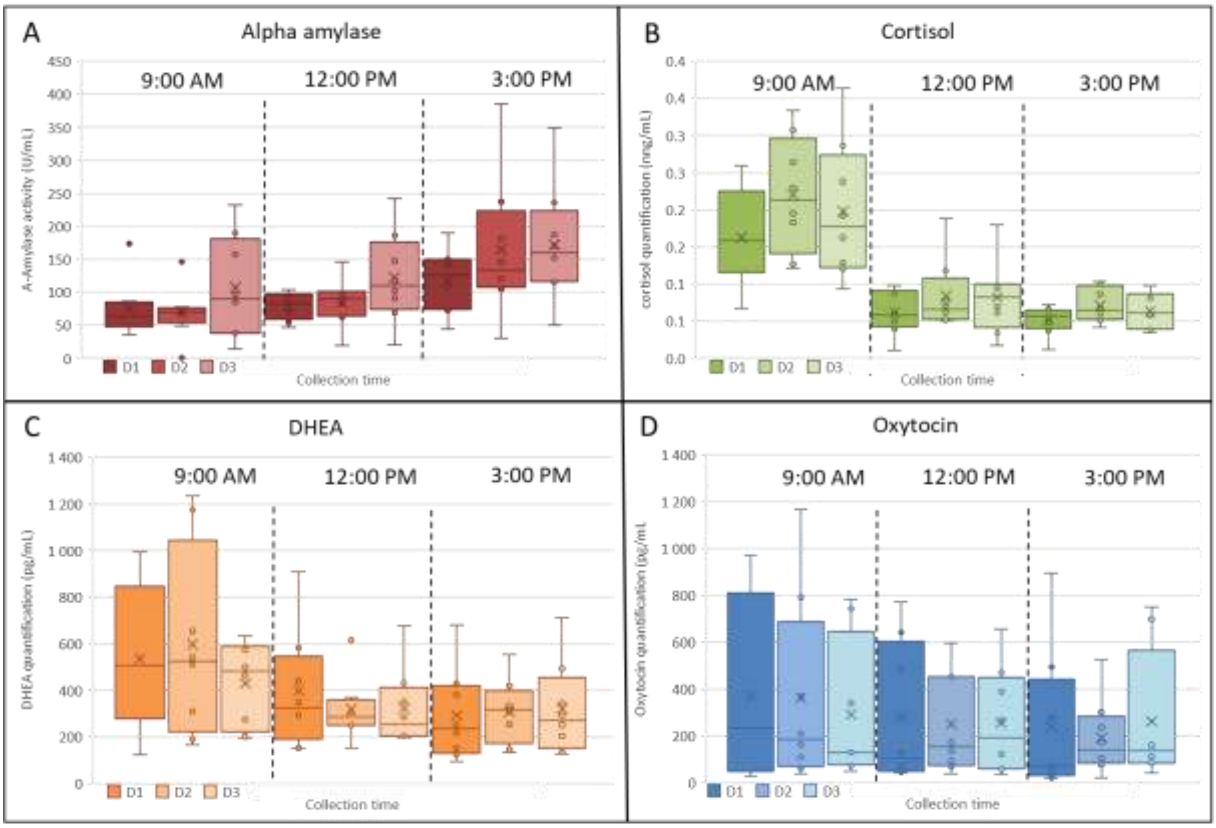


**Fig. 2SM: Salivary biomarker concentrations according to circadian rhythm.** Box plot of biomarker concentrations quantified in 8 salivary samples collected at 3 separate times (9 A.M., 12 P.M., 15 P.M.), 3 consecutive days (D1-D3). **(A)** Salivary alpha-amylase activity (U/mL) ; **(B)** Salivary cortisol concentration (ng/mL) ; **(C)** Salivary DHEA concentration (pg/mL) ; **(D)** Salivary oxytocin concentration (pg/mL). D1: day 1, D2: day 2 and D3: day 3. The median is represented by a line and the mean by a cross.

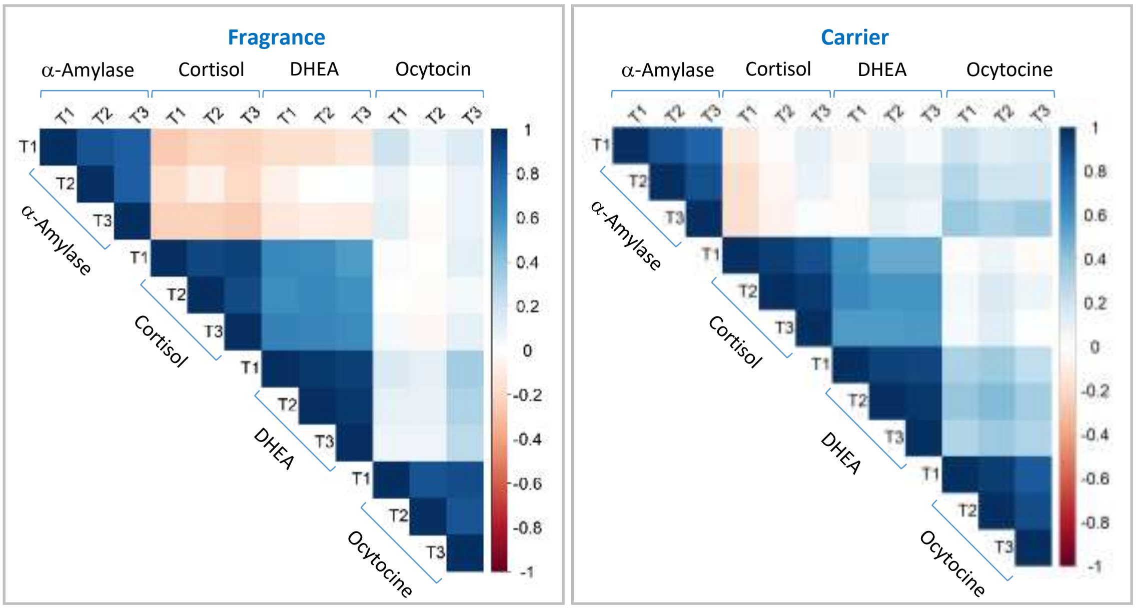


**Fig. 3SM: Correlation of variances between the four biomarkers.** T1: collection of salivary samples 5 min before stimulation; T2: collection of salivary samples 5 min after stimulation; T3: collection of salivary samples 20 min after stimulation. The color scale indicates the degree of correlation between two measurements: 1 = maximum correlation; -1 = maximum anti-correlation. Correlations were calculated using the Spearman method.

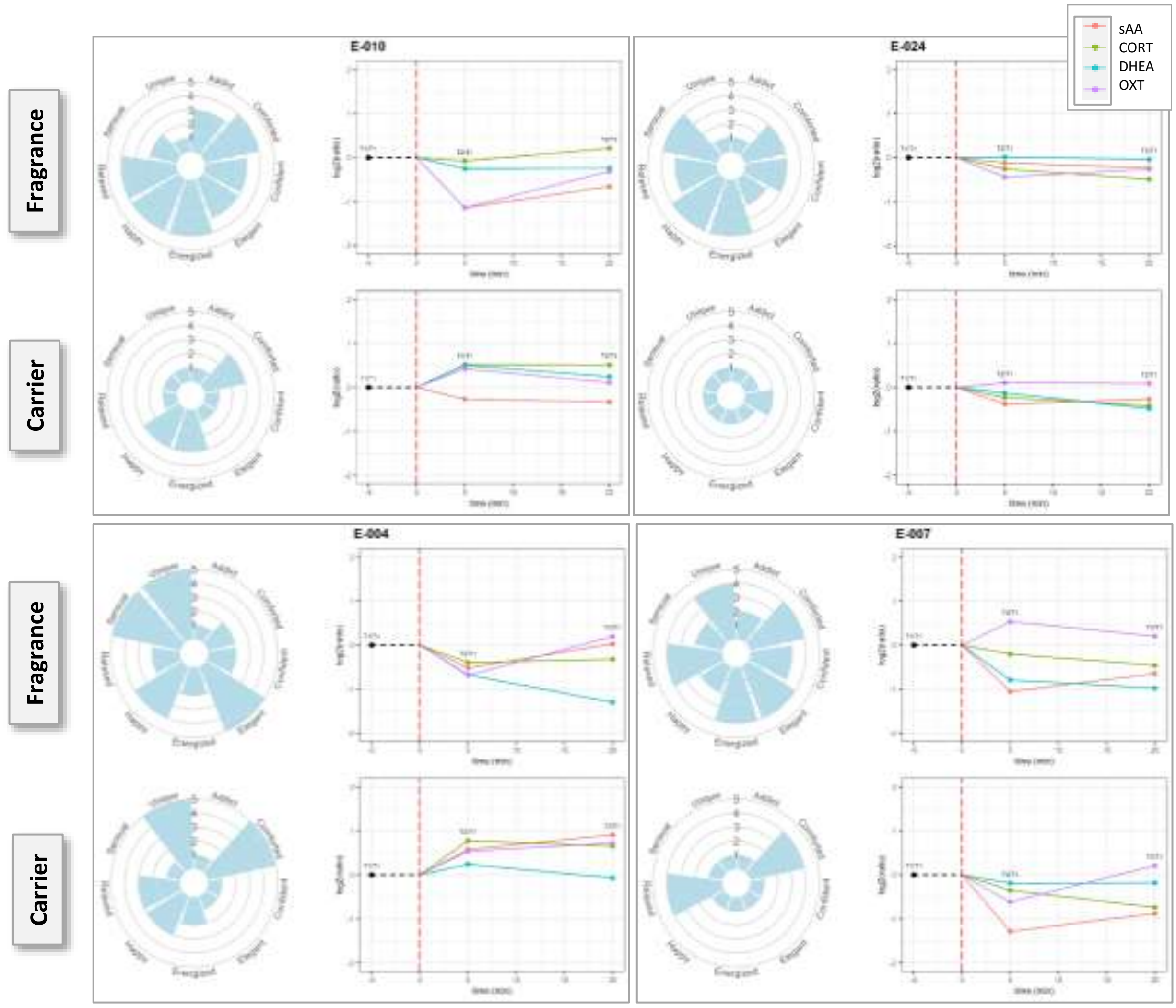


**Fig. 4SM: Example of individual representation of emotions and evolutionary profiles of biomarkers.** The numbers E-0XX correspond to the subjects participating in the study. Radar plots depict, for each participant, the intensity of emotional responses (scale 1–5; left panels): 1 corresponding to the lowest valence and 5 to the highest valence. Biomarker concentrations following exposure to the fragrance (upper panels) or the carrier (lower panels) are shown in the right panel. The time after stimulation (S2 or S3) were compared to the time before stimulation (S1) and represented on a log2 scale. Each curve represents a biomarker.

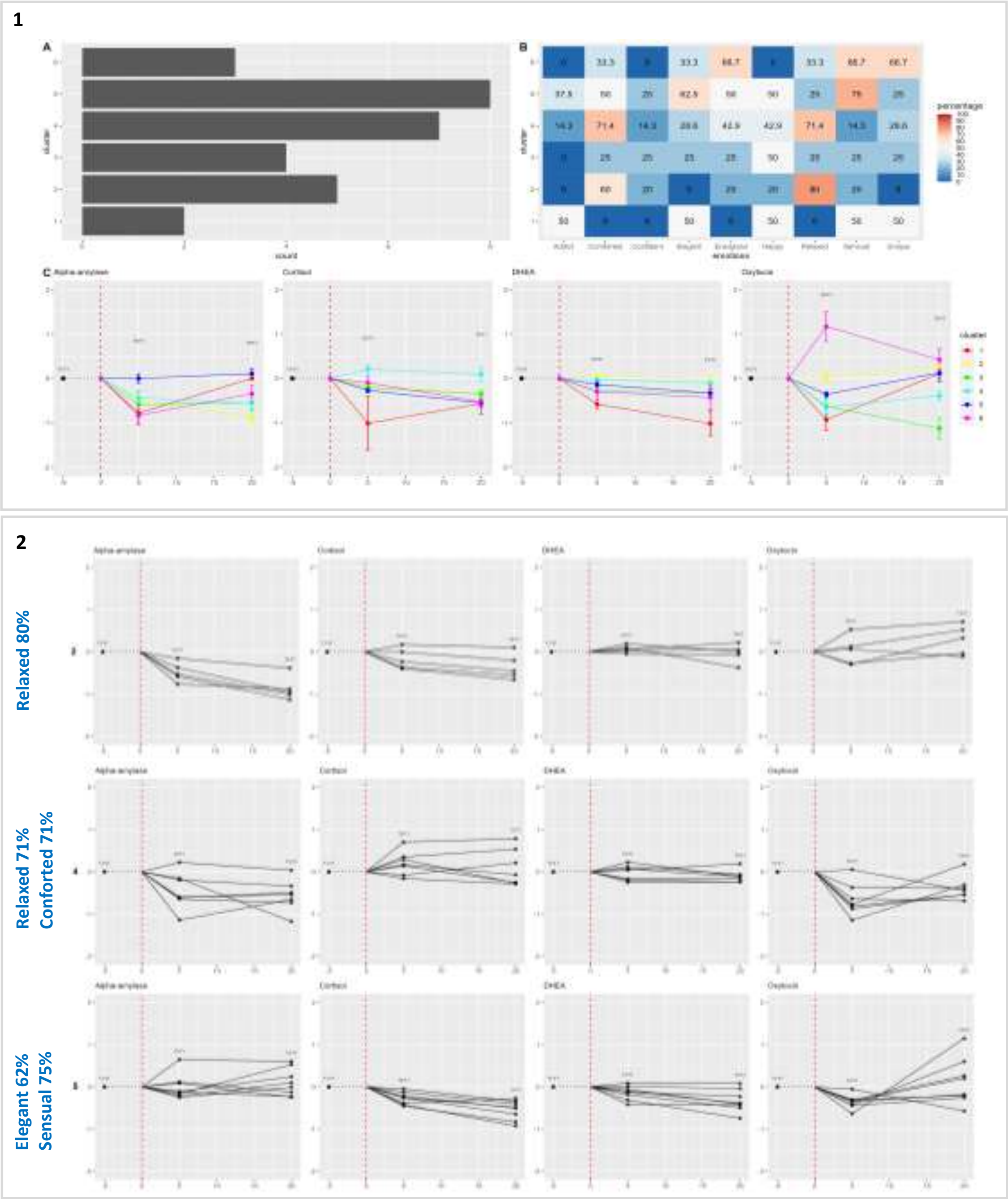


**Fig. 5SM: K-means clustering of individuals based on biomarker profiles into 6 groups. 1)** Global overview of the analysis. **(A)** Number of subject per cluster. **(B)** Heatmap showing the percentage of individuals in each cluster who reported experiencing each emotion. **(C)** Cluster-specific biomarker profiles, with each cluster represented by a different color. Curves show the mean biomarker concentration for individuals in the cluster, with the standard error of the mean also indicated. **2)** Representative individual biomarker profiles for clusters 2, 4 and 5. Data normalization was conducted using the Box-Cox transformation implemented via the MASS library (version 7.3.60.0.1). The visualizations were generated using R software, specifically the ggplot2 (version 3.5.1) and ggpubr (version 0.6.0) libraries.

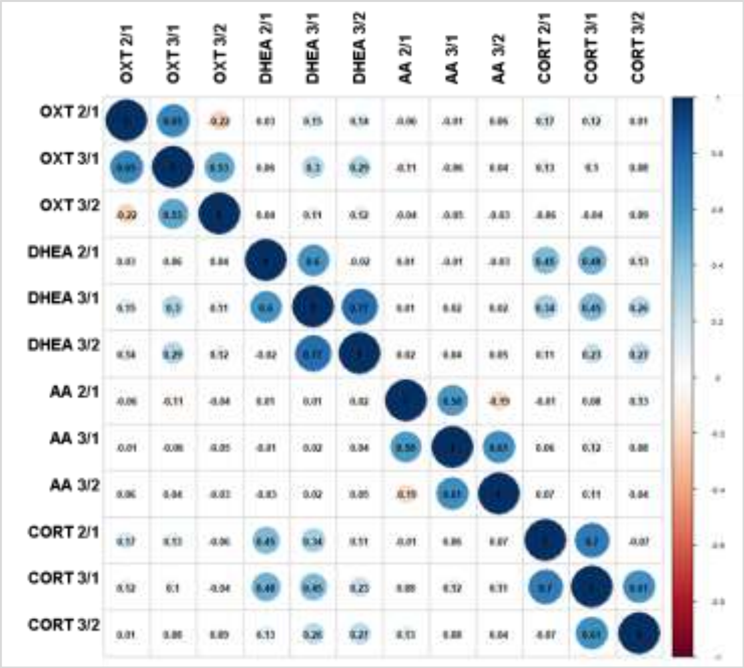


**Fig 6SM: Correlation of salivary biomarker concentration ratios.** The four biomarkers (sAA, CORT, DHEA, OXT) were reliably quantified (mean coefficient of variation <9%) across 708 saliva samples, including 236 unique participant-fragrance combinations with 3 sampling times (1, 2 and 3) each. Correlations were calculated using the Pearson method.

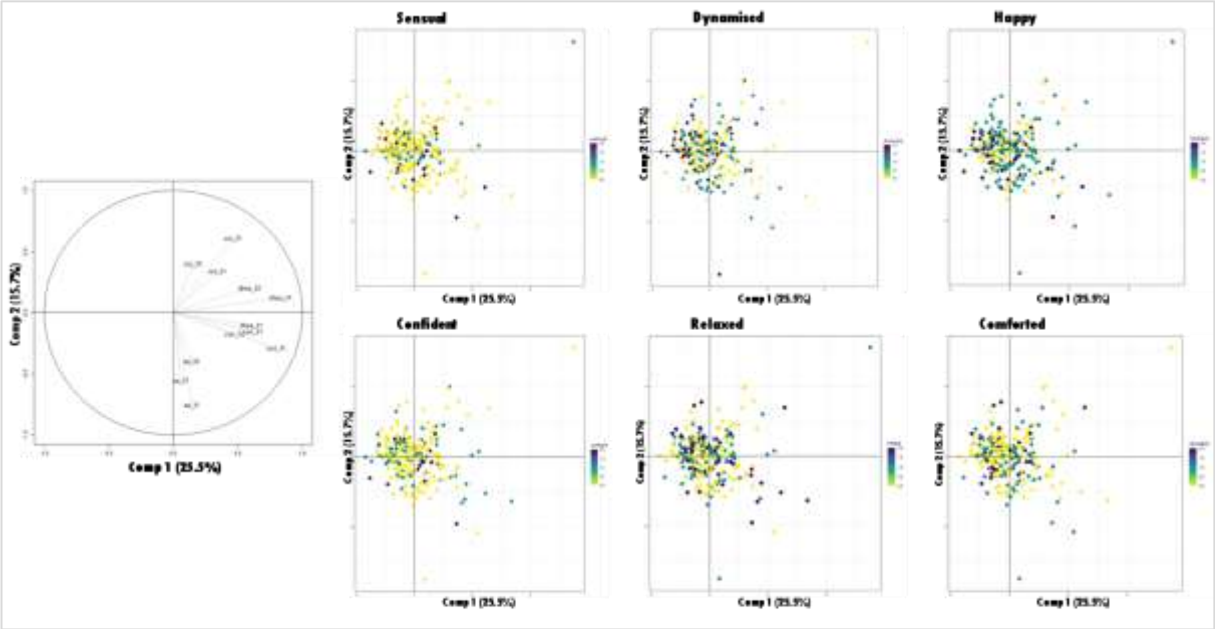


**Fig. 7SM: Principal component analysis (PCA) of biomarkers ratios (S2/S1 and/or S3/S1) in relation to emotions.** PCA were performed using R library factomineR (version 2.11) and factoextra (version 1.0.7), either on the data set from all participants that scented the four fragrances, or by analyzing each fragrance separately. Colors indicated the intensity of emotional responses (scale 0–2): 0 corresponding to the lowest valence (yellow) and 2 to the highest valence (dark blue).

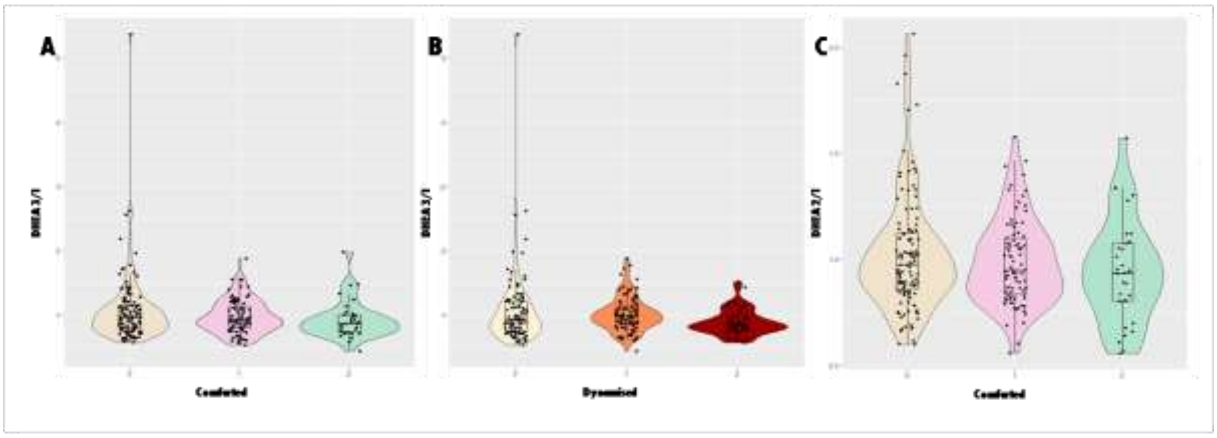


**Fig. 8SM: ANOVA analysis of salivary biomarker ratios among different groups of emotion valence (0 to 2) assessed in the questionnaire.** Groups responding to different levels of emotional valence were compared regarding their biomarker ratios. Emotion valence was assessed on a scale of 0 to 2, with 0 corresponding to the lowest valence and 2 to the highest valence. An ANOVA analysis was performed using each emotion separately or all six emotions together, as continuous or discrete variables, on the entire cohort (59 participants x 4 fragrances = 236 unique combinations). The results presented in the figures above are examples of certain biomarker's ratios that can be predicted by the six emotions combined in a discrete ANOVA analysis when examining at all fragrances. (**A**) DHEA S3/S1 ratio for the comforted emotion; (**B**) DHEA S3/S1 ratio for the dynamized emotion; and (**C**) DHEA S2/S1 for the comforted emotion.

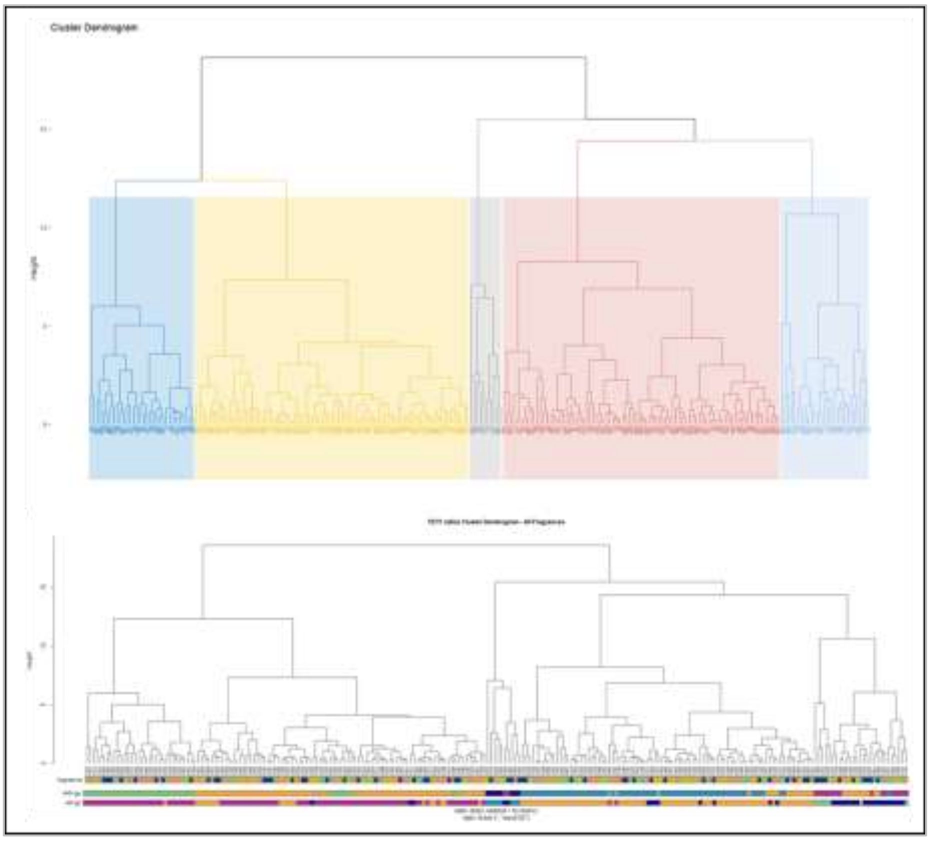

**Fig. 9SM: Participants were classified based on salivary biomarker's ratio profiles using hierarchical k-means (hk-means).** This approach allows to 1) compute hierarchical clustering, 2) cut the tree in k-clusters, 3) compute the center (i.e the mean) of each cluster, 4) do k-means by using the set of cluster centers (defined in step 3) as the initial cluster centers and 5) optimize the clustering. This means that the final optimized partitioning obtained at step 4 might be different from the initial partitioning obtained at step 2. This example shows the hk-means clustering performed on the total cohort (59 participants x 4 fragrances: 236 unique combinations) using the biomarker ratios S2/S1. Participants were nested into five clusters composed of 69, 46, 18, 95 and 8 individuals.

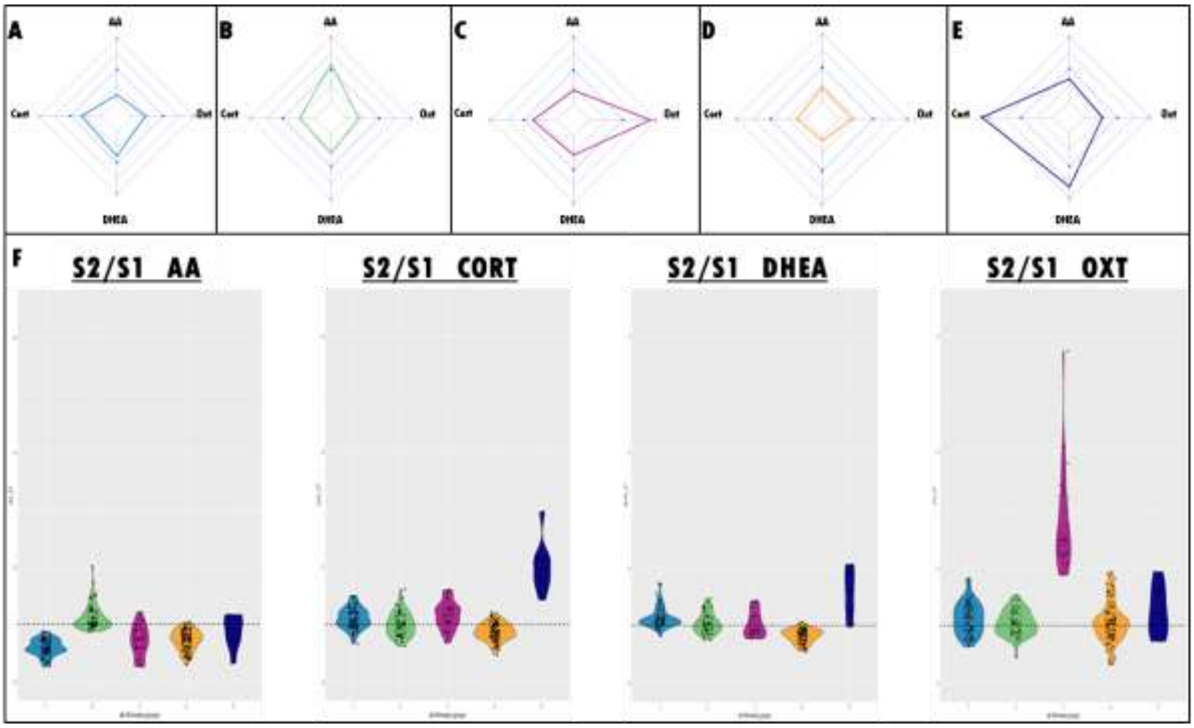


**Fig. 10SM: Salivary biomarker profiles within clusters identified by hk-means**. Salivary biomarker ratio (S2/S1) profiles for each cluster nested by hk-means performed on the total cohort (59 participants x 4 fragrances: 236 unique combinations). Spiderplots show the absolute ratio values for each cluster; **(A)** cluster 1: 69 individuals; **(B)** cluster 2: 46 individuals; **(C)** cluster 3: 18 individuals, **(D)** cluster 4: 95 individuals and **(E)** cluster 5: 8 individuals. **(F)** Comparison of the distribution of biomarker ratios among clusters represented in violin plots. Light blue: cluster 1; green: cluster 2; purple: cluster 3; orange: cluster 4 and dark blue: cluster 5.

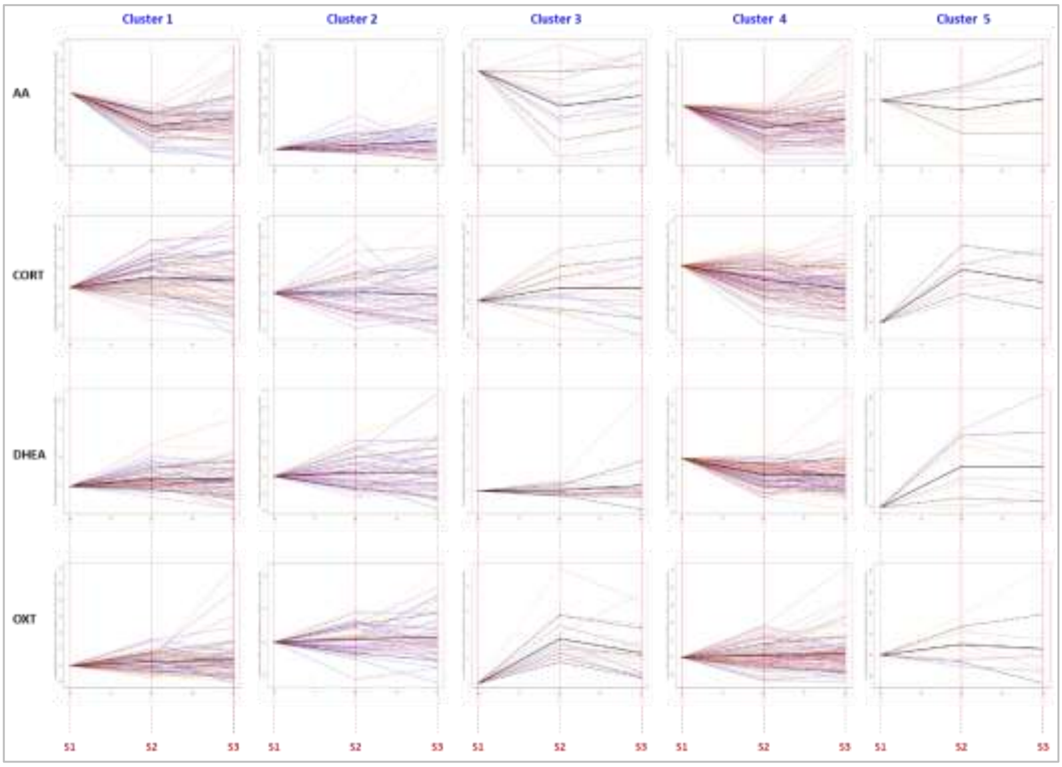


**Fig. 11SM: Temporal evolution of individual salivary biomarker ratios (S1/S1; S2/S1; S3/S1) in clusters identified by hk-means.** The analysis was performed on the total cohort (59 participants x 4 fragrances: 236 unique combinations). Curves represent individual biomarkers' ratios in different sampling (S1, S2 and S3) normalized to S1 biomarkers' concentrations of each participant. Curves are colored by fragrance and the mean of each cluster for each biomarker is represented by the bold black line. Standard deviation is represented by dotted black lines.

**Table 3SM: Questionnaire-based emotional profiles in hk-means clusters.** Differential emotional responses were evaluated between clusters identified by hk-means performed either on the whole cohort (236 unique combinations participants/fragrances) or by fragrance separately using Chi2 test. Emotional responses were discretized (questionnaire response O = negative response to a given emotion; questionnaire response 1 and 2 were nested together = positive response to a given emotion). Data of the p-values are shown in the table.

| | **Chi2 *p*-values** | | | | | |
|---|---|---|---|---|---|---|
| | Dynamised | Happy | Relaxed | Comforted | Sensual | Confident |
| **ALL** | >0.5 | >0.5 | >0.5 | >0.5 | >0.5 | **0.23** |
| **J3** | **0.3** | >0.5 | **0.31** | >0.5 | >0.5 | **0.23** |
| **J4** | >0.5 | **0.2** | >0.5 | **0.34** | >0.5 | **0.23** |
| **J5** | >0.5 | >0.5 | >0.5 | >0.5 | >0.5 | **0.29** |
| **J6** | >0.5 | >0.5 | >0.5 | **0.08** | **0.14** | **0.27** |

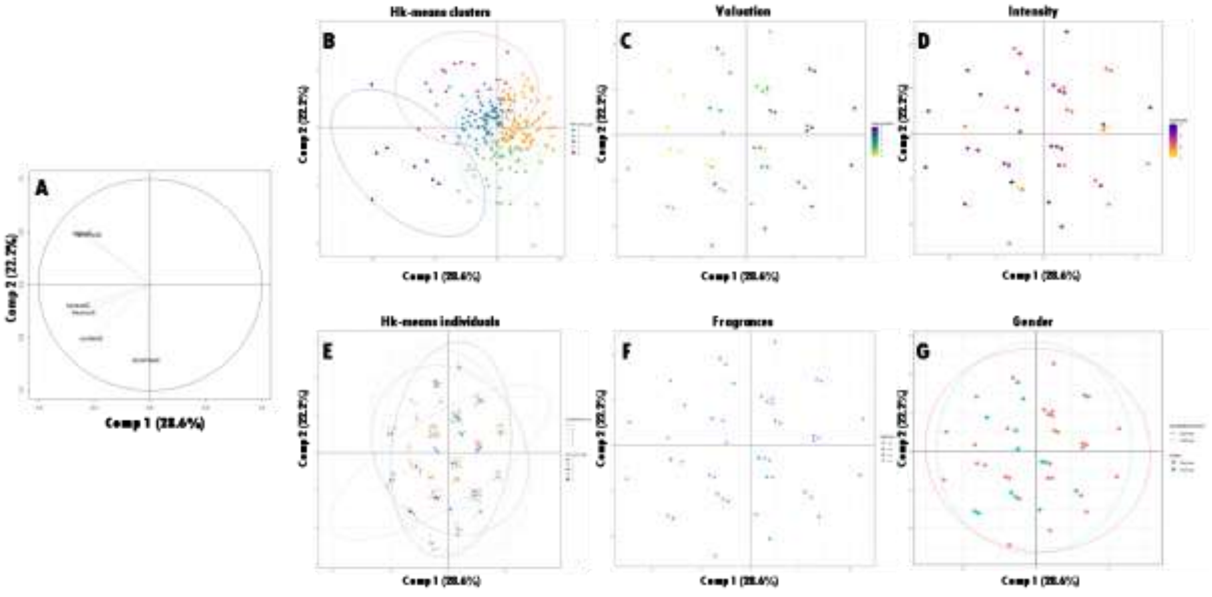


**Fig. 12SM: Principal component analysis (PCA) was performed on discretized binary emotions and correlations to different variables were analyzed (B – G). (A)** The emotion responses used as quantitative variables were discretized: questionnaire response O = negative response to a given emotion; questionnaire response 1 and 2 were nested together = positive response to a given emotion. PCA was performed using R library factomineR (version 2.11) and factoextra (version 1.0.7), on the data from the whole cohort (236 unique combinations) with the four fragrances and the clusters projected were nested using the hk-means clusterization. No pattern distribution was observed for any of the analyzed variable: **(B)** Hk-means clusters groups; **(C)** Valuation; **(D)** Intensity; **(E)** Same as **(B)** with some random effects to visualize the presence of multiples measures with identical coordinates due to the limited number of variables (6 emotions) with only two possible values (0; 1). Large point correspond to true values; smaller points are randomly distributed around their real values; **(F)** Fragrances; **(G)** Gender. A similar analysis was also performed using hk-means clusterization performed by fragrance separately.

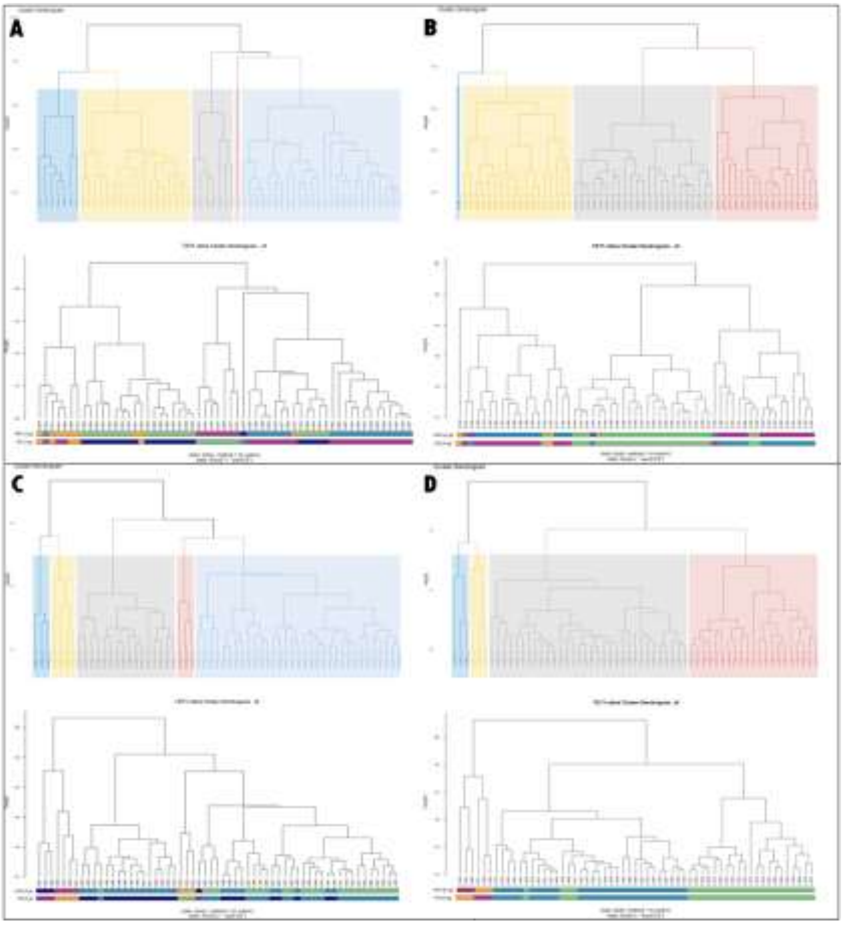


**Fig. 13SM: Classification of participants based their salivary biomarker ratios (S2/S1) for each independent fragrance using hierarchical k-means (hk-means).** **(A)** The fragrance J3 nested into five clusters consisting of 21, 22, 7, 9 and 1 individuals, respectively. **(B)** The fragrance J4 nested into four clusters composed of 16, 25, 16 and 2 individuals, respectively. **(C)** The fragrance J5 nested into five clusters composed of 24, 24, 4, 3, and 4 individuals, respectively. **(D)** The fragrance J6 nested into four clusters composed of 28, 25, 3 and 3 individuals, respectively.

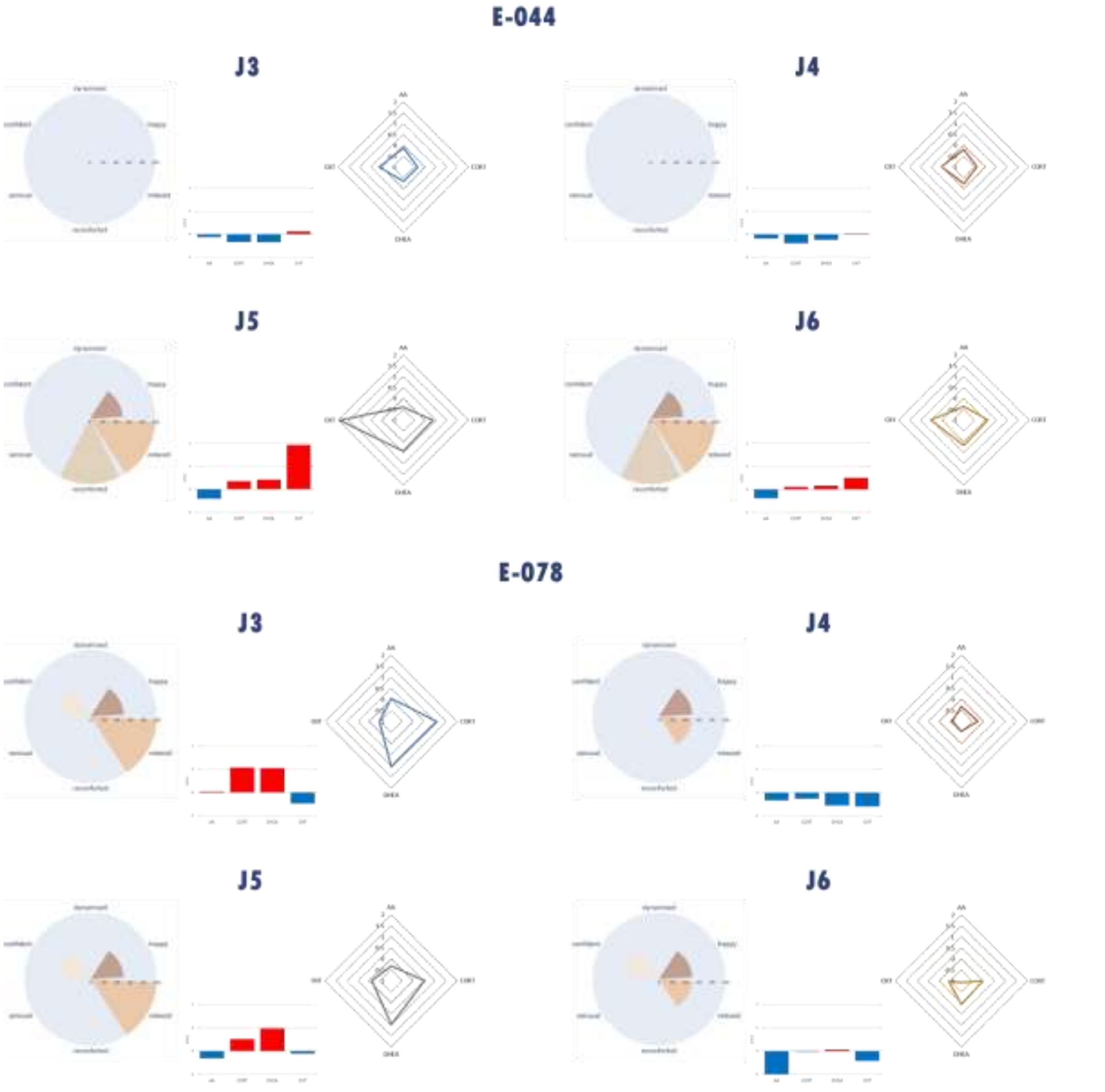


**Fig. 14SM: Individual case studies of emotional responses and biomarker ratio (S2/S1) profiles by fragrance.** For each fragrance (J3–J6), clusters are shown with emotional response profiles (left, radar plots; % participants per cluster) and log2 mean biomarker ratios (S2/S1) (center, histograms; right, spider plots). Participant E-044 is shown on the upper panel and participant E-078 on the lower panel. The visualizations were created using Python libraries matplotlib (version 3.7.4), seaborn (version 0.13.2), and plotly (version 5.9.0).

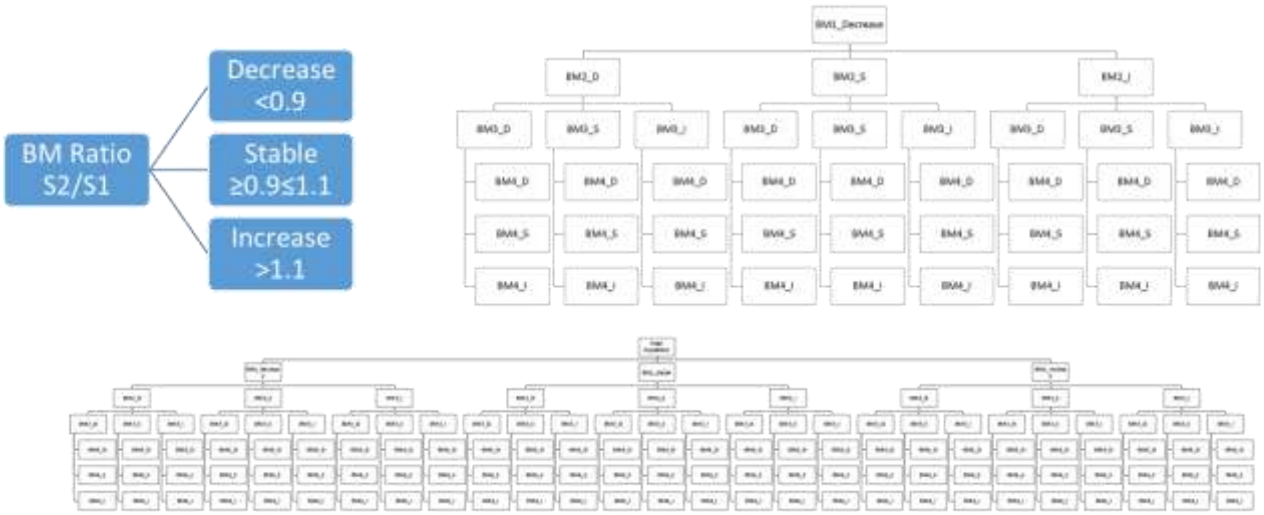


**Fig. 15SM: Discretization of salivary biomarker ratios (S2/S1) for new clustering approaches.** Biomarkers ratios were discretized into three categories: i.e. <0.9 = decrease; ≥0.9≤1.1 = stable and >1.1 = increase. By combining the discretized biomarker ratios (e.g. category 1 = alpha-amylase decrease, cortisol decrease, DHEA decrease, oxytocin decrease; category 2 = alpha-amylase decrease, cortisol decrease, DHEA decrease, oxytocin increase, and so on), 81 putative categories were defined.

| | Dynamised | Happy | Confident | Comforted | Relaxed |
|---|---|---|---|---|---|
| True positive | 104 | 162 | 20 | 50 | 103 |
| Total positive | 135 | 167 | 84 | 115 | 136 |
| **Sensitivity** | **77%** | **97%** | **24%** | **43%** | **76%** |
| True negative | 52 | 9 | 139 | 99 | 45 |
| Total negative | 101 | 69 | 152 | 121 | 100 |
| **Specificity** | **51%** | **13%** | **91%** | **82%** | **45%** |

**Table 4SM: Performances of salivary biomarker patterns for emotion identification**. The biomarkers-based emotional responses were compared to the questionnaire-based emotional responses for each participant, all fragrance combined. For each emotion, we calculated a sensibility (% True positives/Total positives) and a specificity (% True negatives/Total negatives). Biomarker patterns used in this approach were those identified from the decision trees generated by CART performed using the entire cohort data (236 unique combinations of individuals/fragrances) and biomarkers' ratios S2/S1 as discrete variables.

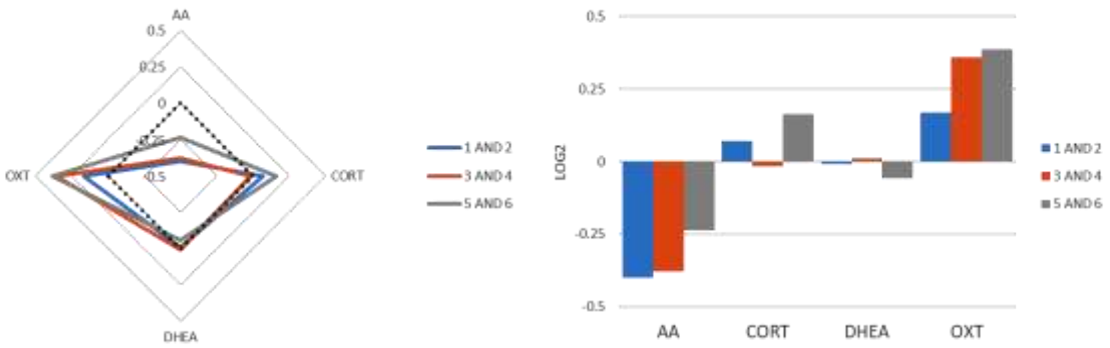


**Fig. 16SM: Salivary oxytocin and alpha-amylase ratio (S2/S1) levels associate with valuation of all fragrances together.** In the questionnaire, each participant rated their valuation for each fragrance on a scale of 1 to 6: 5 /6 corresponding to lower valuation; 3/4 to medium valuation and 1/2 higher valuation. Radar plot describes the average levels of the ratio (S2/S1) for each biomarker according to the valuation groups (left panel). On right panel, histograms show the mean ratios of alpha-amylase and oxytocin ratios according to the fragrance valuation groups. This analysis was performed using data from the entire cohort (236 unique combinations of participants/fragrances).
